# Endocrinology shadows ecology: Characterization of Markhor (*Capra falconeri heptneri*) reproductive cycles through non-invasive hormone assessment

**DOI:** 10.64898/2026.04.20.719681

**Authors:** Bharti Arora, Shiwangi Rai, Pranita Gupta, Joy Dey, Basavaraj S Holeyachi, Samrat Mondol

**Affiliations:** Wildlife Institute of India, Chandrabani, Dehradun, 248001, India; Mammal Research Institute, Department of Zoology and Entomology, University of Pretoria, 0028, South Africa; Padmaja Naidu Himalayan Zoological Park, Darjeeling, West Bengal, India; Academy of Scientific and Innovative Research (AcSIR), Ghaziabad-201002, India

**Author notes:** Corresponding author: Samrat Mondol, PhD. Wildlife Institute of India, Chandrabani, Dehradun, India.

**Keywords:** Reproductive function, Longitudinal monitoring, Pregnancy detection, Nonreproductive phase, Conservation breeding, Animal welfare

## Abstract

Markhor (*Capra falconeri*) is a charismatic, threatened, large, high-altitude bovid found in parts of central and south Asia. The species faces threats such as habitat loss, hunting, poaching, livestock competition, hybridisation, and disease, yet research on wild populations is challenging. Various biological aspects, including surveys, diet, population dynamics, interactions with livestock, hybridisation, and disease, have been studied locally, along with behavior and reproductive biology, but details such as pregnancy, oestrus, and parturition timing remain unestablished. We conducted the first systematic, detailed, and fine-scale characterization of the reproductive steroid profiles of two males and five female markhors (*Capra falconeri heptneri*) in a captive population at the Padmaja Naidu Himalayan Zoological Park (PNHZP), West Bengal, India. We collected weekly fecal samples, standardized and validated measurements of progesterone (fP4M) and testosterone (fTM) metabolites, and conducted reproductive profiling to assess reproductive stages in both sexes. Analyses of annual fP4M and fTM data from male and female markhor individuals showed similar profiles and synchronicity, with individual variation, and peaks and baselines were evident for both hormones. In both sexes, significantly higher hormone titres were observed during the sexually active and inactive phases. Non-invasive measurement of reproductive hormones accurately reflected ovarian function in females, helping establish mating, gestation, and parturition timelines in female markhors and determine the breeding season in males. These approaches support husbandry and breeding management by identifying optimal pairing, diagnosing pregnancy, and predicting parturition in both captive and wild populations. When applied correctly, these tools could greatly aid population monitoring of other endangered species across high-altitude regions worldwide.

## Introduction

Markhor (*Capra falconeri*) is one of the most charismatic high-altitude bovids and the largest member of the genus *Capra* (Michel et al., 2015). They are found exclusively in the semi-arid and mountainous regions of north-eastern Afghanistan, northern India (southwest Jammu and Kashmir), northern and central Pakistan, southern Tajikistan, south-western Turkmenistan, and southern Uzbekistan (Grubb, 2005). Based on their geographical distribution and distinctive horn morphology (flared versus straight horns), they are further classified into three subspecies: *Capra falconeri falconeri* (flared-horned subspecies found across Afghanistan, Pakistan, and India), *Capra falconeri heptneri* (flared-horned subspecies found in Afghanistan, Tajikistan, Uzbekistan, and Turkmenistan) (Grubb, 2005), and *Capra falconeri megaceros* (straight-horned subspecies found in Afghanistan, Pakistan) (Michel et al., 2015). Across their range, the species faces severe human-related impacts, including habitat loss and fragmentation, traditional hunting, poaching, competition from livestock grazing, hybridisation, and disease transmission. Despite these challenges, significant conservation efforts, including the creation and preservation of protected areas and tangible community conservation initiatives, have led to the recovery of local populations in many regions. With a global population of fewer than 6000 mature individuals and an increasing population trend, the species is classified as ‘Near Threatened’ (criterion C2a(i)) by the IUCN (Michel et al., 2015), Appendix I of CITES, and Schedule I of the Indian Wildlife Protection Act 1972.

Research on wild Markhor populations is challenging due to their rugged, high-altitude habitats and cross-border presence across different geopolitical regions (Bhatnagar et al., 2009). However, various aspects of Markhor biology, including population status surveys (Roberts and Bernhard, 1977), diet (Bashir et al., 2020), population dynamics (Ali, 2008), interactions with livestock (Karlstetter, 2008), hybridisation status (Hammer et al., 2008), and disease outbreaks (Peyraud et al., 2014; Woodford et al., 2004), have been studied at local levels.

Significant effort has been dedicated to ex situ conservation, establishing captive populations across North America, Europe, and Asia, including India (Asif, 2019; Shackleton, 1997). Notably, *Capra falconeri heptneri* has a sizable captive population of around 400 individuals housed in zoological parks and semi-open habitats such as the Lesser Carpathians in Slovakia, under the ARES markhor breeding programme (Pokoradi, 2005). These initiatives have addressed various aspects of the species’ biology, such as behaviour (Ali, 2008; Roberts and Bernhard, 1977) and male reproductive biology (Bezjian et al., 2013). However, pregnancy assessment, timing of estrus, parturition, and duration of sexual activity have not been established in the Markhor. In the wild, various reproductive behaviours such as rut, gestation, and parturition have been documented through visual observations (Huffman, 2003; Michel and Michel, 2015; Roberts, 1977), but no physiological data are available. Due to the limited detailed information available on this species’ reproductive biology, further research is necessary to support breeding programmes and management strategies aimed at the long-term conservation of the species.

In this study, we conducted a systematic, detailed, and fine-scale characterisation of the reproductive steroid profiles of both male and female Markhor (*Capra falconeri heptneri*) in a captive population at the Padmaja Naidu Himalayan Zoological Park (PNHZP), West Bengal, India. This is the only zoological park in India that hosts a captive Markhor population and actively participates in their conservation breeding programmes. The primary objectives of this study were: (a) the biochemical and biological validation of female (Progesterone or P4) and male (Testosterone or T) hormones from non-invasive samples (fecal pellets); (b) to generate fecal hormone profiles from multiple individuals to evaluate variations in reproductive hormone levels; and (c) to estimate different reproductive phases in Markhor. The standardized methods and baseline information would offer valuable support for monitoring the success of various assisted reproductive techniques (ARTs) in captive facilities and for reliably assessing reproductive parameters in markhor across its range.

## 2. Materials and methods

### 2.1 Research permissions and ethical considerations

The work was conducted as part of the study titled “Study on behaviour and reproductive biology of Markhor (*Capra falconeri*) at PNHZ Park”, approved by the West Bengal Zoo Authority (Letter no. 104/WBZA/T-18(j)/21-22 dated 15/06/2021). No ethical clearance was required because the study used non-invasive samples.

### 2.2 Study area

The study was conducted at the Padmaja Naidu Himalayan Zoological Park (hereinafter referred to as PNHZP) in the Darjeeling district of West Bengal, India. The zoo is recognized by the Central Zoo Authority, Government of India, as a highly specialized, medium-sized, high-altitude facility, situated at an altitude of 2,137 meters. The zoo spans across 67.57 acres and houses a diverse range of carnivores, herbivores, and omnivores. PNHZP is the national coordinating zoo for several high-altitude species conservation breeding programs, including the Red Panda (*Ailurus fulgens*), Himalayan Salamander (*Tylototriton himalayanus*), Himalayan Wolf (*Canis lupus himalayanus*), Satyr Tragopan (*Tragopan satyra*), and Snow Leopard (*Panthera uncia*). It also participates in the conservation of other high-altitude specialist species such as Blue Sheep (*Pseudois nayaur*), Bhutan Grey Peacock Pheasant (*Polyplectron bicalcaratum*), Himalayan Tahr (*Hemitragus jemlahicus*), Himalayan Blood Pheasant (*Ithaginis cruentus*), and Himalayan Monal (*Lophophorus impejanus*).

### 2.3 Study subjects and housing

We selected five markhor individuals (*Capra falconeri heptneri*) at PNHZP for this longitudinal study: three females and two males (Table 1). Among them, one male (Arvind, 15 years) and one female (Sonia, 10 years) were brought from the Augsburg Zoo in Germany. The other two females (Ivy: 6 years and Supriya: 5 years) and one male (Ranbir: 10 years) were born in captivity and were all unrelated.

**Table 1:**
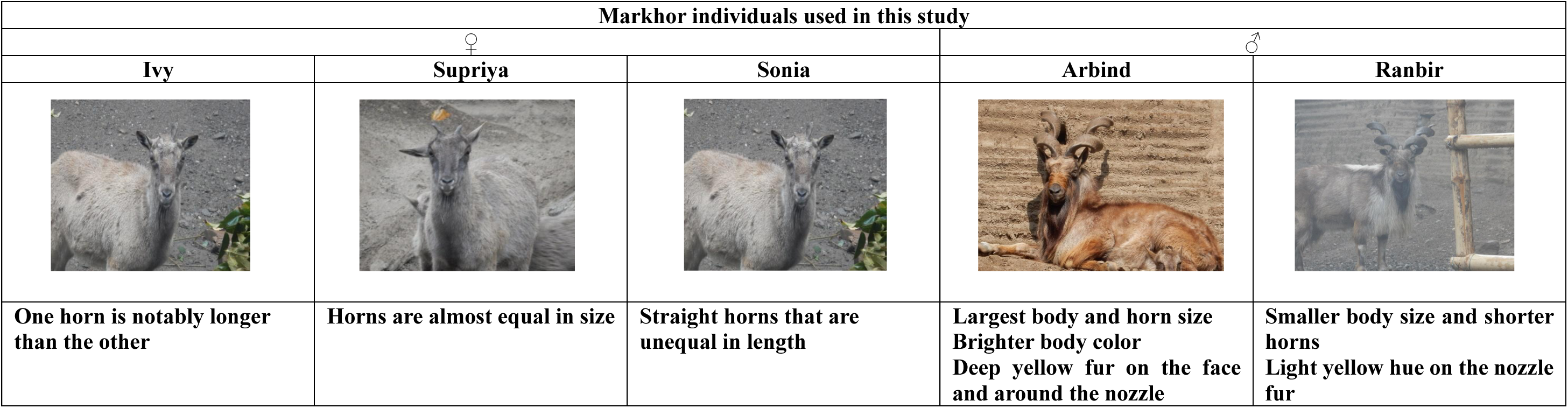
The identifying details of the markhor study subjects in PNHZP.

| Markhor individuals used in this study |  |  |  |  |
| --- | --- | --- | --- | --- |
| ♀ |  |  | ♂ |  |
| Ivy | Supriya | Sonia | Arbind | Ranbir |
| One horn is notably longer than the other | Horns are almost equal in size | Straight horns that are unequal in length | Largest body and horn size<br>Brighter body color<br>Deep yellow fur on the face and around the nozzle | Smaller body size and shorter horns<br>Light yellow hue on the nozzle fur |

The study subjects were housed together in an enclosure measuring 550 m² (26.35 × 20.90 m) (Supplementary Figure 1). The animals remained in the same configurations throughout the study duration, allowing them to become accustomed to the enclosure and management practices. The enclosure provided the subjects with a mix of natural elements, such as trees, and man-made structures (i.e., shelters, feeding boxes, and platforms), along with an open, shaded area that featured a designated water area to promote the overall well-being of the species. All individuals had free access to the exhibit.

A dedicated team comprising one veterinary officer, one zoo biologist, and two animal keepers performed all husbandry tasks for the subjects. The keepers provided twice-daily feeding to all animals in the exhibit (Supplementary Table 1). The enclosure was cleaned daily before morning feeding hours. The zoo veterinary team conducted periodic health examinations to ensure routine health management for all these animals.

### 2.4 Experimental Design

We conducted a longitudinal fecal sampling from May 2022 to April 2023 in each study subject to characterize the sex-specific reproductive cycles. In the enclosure, the animal keepers and researchers identified each study subject by unique morphological attributes (e.g., horn shape, size, body size, color) (Table 1) and closely monitored individual reproductive behaviors during the study period. We adopted a focal-animal sampling approach for each identified individual. We collected 1-2 samples/individual/week for males, whereas 2-3 samples/individual/week were collected for females, totaling 206 samples (♂n = 63, ♀n = 143). Relatively fresh fecal pellets (less than six hours old) were collected from each captive individual following Biswas et al. (2019) and were stored in a −20 °C freezer at the zoo. Monthly sample batches were shipped through cold chains to the Wildlife Endocrinology Facility at the Wildlife Institute of India, Dehradun, India, where they were stored at −20 °C until processing.

### 2.5 Fecal hormone extraction and assays

During sample processing, the frozen fecal samples were thawed overnight, pulverized, and oven-dried in a Promax Heating oven (Techno Engineer, India) at 50 °C for 48 hours (Patel et al., 2021; Patel et al., 2023). The dried samples were sifted through separate 1-mm mesh, and the fecal powders were collected and homogenized. Furthermore, fecal hormone metabolites were extracted from well-mixed 100 mg of each powder using 15 mL of 70% ethanol, following the protocols described in (Patel et al., 2021; Patel et al., 2023). After completion, 4 mL of each extract was stored in cryochill vials at −20 °C before further assays.

We measured two reproductive hormone metabolites from the field-collected samples: Progesterone (fP4M) and Testosterone (fTM), using commercially available EIA kits (P4-K025-H5 and T-K032-H5, Arbor Assays, Ann Arbor, USA). The assay cross-reactivities are listed in Table 2. Sample extracts were incubator-dried (Promax Heating oven, Techno Engineer, India) at 30 °C and then resuspended in assay buffers. All assays were performed in duplicate according to the respective manufacturer’s protocols. Hormone concentrations were expressed as µg/g of fecal powder.

**Table 2:** Details of the hormone EIAs conducted in this study.

| Hormone | Dilution | Parallelism | Accuracy |  |  | Inter-assay CV | Intra-assay CV | Cross-reactivities |
| --- | --- | --- | --- | --- | --- | --- | --- | --- |
|  |  |  | Slope | R<sup>2</sup> | Linear equation |  |  |  |
| Testosterone (fTM) | 1:120 | F<sub>1,7</sub> = 0.149<br>P = 0.71 | 0.9133 | 0.9894 | Y=0.9133x+0.0198 | 8.9% | 3.3% | Testosterone 100%, 5α Dihydrotestosterone 56.8%, 11 Ketotestosterone 2.34%, Androstendione <1%, Androsterone <1%, DHEA, Cholesterol <1%, 17β Estradiol <1%, Progesterone <1%, Pregnenolone <1%, Hydrocortisone <1%, Cholic Acid <1%, Cholic Derivatives <1%, Aldosterone <1%, Cortisol 1%, Corticosterone 1%, Cortisone 1% |
| Progesterone (fP4M) | 1:240 | F<sub>1,6</sub> = 3.175<br>P = 0.12 | 0.9857 | 0.9956 | Y=0.9857x+0.0189 | 6.7% | 4.8% | Progesterone 100%, 3β- hydroxy-Progesterone 172%, 3α-hydroxy progesterone 188%, 11α-hydroxy progesterone 147%, 11β hydroxyprogesterone 2.7%, 5-α-hydroprogesterone 7%, Pregnenolone 5.9%, Corticosterone <1%, Androstenedione <1% |

### 2.6 Assay validations

We conducted biochemical (parallelism and accuracy) and biological validations to assess the assay performance for both hormones. Parallelism was evaluated to determine reliable hormone quantification across concentrations and to identify optimal dilutions (at 50% binding), using serial dilutions (1:15, 1:30, 1:60, 1:120, 1:240, 1:480, 1:960) of pooled markhor fecal extracts (n=15; three samples from each study subject). The output of the sigmoidal response curve was statistically tested against the standard-calibrator response curve of each hormone. The parallelism results were examined using an F-ratio test for differences in slopes. Furthermore, fP4M and fTM accuracy tests were performed to assess potential interference during antibody interactions by spiking hormone extracts with equal volumes of diluted fecal extracts and assaying with standards. The values were graphed as regression lines to determine the observed and expected concentrations, with a slope of 1 (ranging from 0.9 to 1.1) (Keel, 1996). Table 2 demonstrates the dilutions used for the assays.

For biological validation of P4 and T measurements, we used female fecal pellets immediately after birth and male pellets following reproductive behaviors (for example, mounting). We compared these with samples from the same individuals when reproductive behavior was not observed.

### 2.7 Estimation of different reproductive phases in markhors

The systematic weekly sampling of known individuals of both sexes at the zoo enabled us to investigate the temporal patterns of reproductive events, such as gestation duration and mating period. P4 is essential for the establishment and maintenance of mammalian pregnancies (Schuler et al., 2018; Spencer et al., 2004). Pregnancy onset is characterized by a prolonged increase in physiological progesterone levels, whereas progesterone levels drop significantly during parturition (Smith et al., 2009). Multiple mounting and copulation attempts coincide with rising T levels (Mokkonen et al., 2012). We employed an iterative process to establish a baseline (Brown et al., 1996) using the *hormLong* package (Fanson and Fanson, 2015) in R v4.2.2 to correlate quantified hormone metabolite concentrations associated with different reproductive events in Markhor. fP4M and fTM concentrations exceeding the mean (M) plus 1.5 times the standard deviation (SD) were discarded, as this would detect smaller hormonal peaks in all individuals (Fanson and Fanson, 2015). The M + 1.5SD iteration was recalculated until no value exceeded this criterion (LaDue et al., 2022). The remaining fP4M and fTM values defined the baseline concentration and established correlations with temporal reproductive events. Concentration values ≥ M + 1.5 SD were considered peaks, indicating significantly higher concentrations. Therefore, a sustained increase in fP4M above baseline defined the onset of pregnancy, while a continuous rise in fTM in males indicated sexual behavior and spermatogenesis (Negro et al., 2010).

### 2.8 Statistical Analyses

We categorized individual Markhor fecal hormone data monthly to investigate reproductive episodes, including mating, pregnancy, gestation, and parturition. Each data point was log-transformed, and Q-Q residual plots and Shapiro-Wilk tests were conducted using Prism version 9.3 (GraphPad; San Diego, CA) to assess normality. Because the data were not normally distributed, we used nonparametric tests in PAST 5.0 (Paleontological Statistics Software, Oslo, Norway) for further analyses. The Kruskal-Wallis test and post-hoc Dunn’s method were employed for multiple group comparisons (monthly median concentrations of fP4M and fTM), while the Mann-Whitney test was used for two-group comparisons (pregnant and non-pregnant months in females; sexual activity and inactivity months in males) using the mean concentrations. The p-value for hypothesis testing was set to α=0.05 for all analyses. All the tests performed were two-tailed.

## 3. Results

### 3.1 Standardization of the target hormone metabolites

Both fP4M and fTM parallelism and accuracy tests demonstrated reliable measurements from Markhor fecal samples across various concentration ranges. During the parallelism test, no significant difference was observed between the binding curves of the diluted samples and the hormone standards (P > 0.05) (Table 2 and Fig. 1). The accuracy tests yielded reliable slopes (Table 2 and Fig. 1), indicating that the fecal extracts did not interfere with the precision of metabolite measurements. The intra-assay coefficient of variation (CV) for P4 and T was 4.8% and 3.3% while the inter-assay CV was 8.9% and 6.7%, respectively (Table 2).

**Figure 1:**
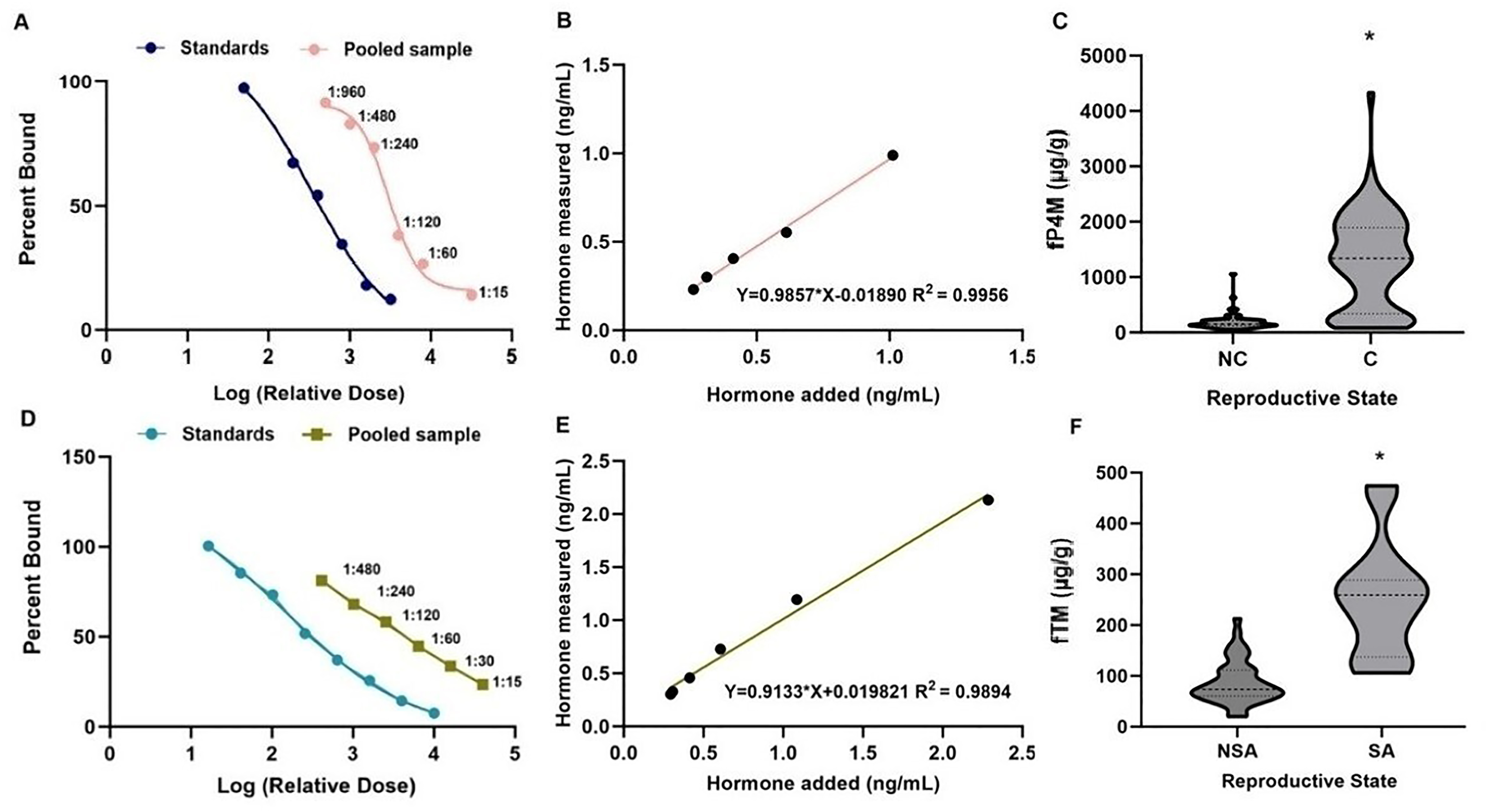
Standardization and validation of fecal hormone metabolites in captive markhors. The top panel shows parallelism (A), accuracy (B) and biological validation (C) for progesterone (fP4M), whereas the bottom panel shows parallelism (D), accuracy (E) and biological validation (F) for testosterone (fTM). * indicates statistical significance between the two groups (p < 0.005).

Biological validation comparisons showed the expected outcomes, with pregnant female fP4M (1340 µg/g) measures significantly higher than those at the non-pregnant stage (152.5 µg/g) (U = 0, p < 0.0001). Sexually active males (259.2 µg/g) also showed significantly higher fTM titres than during the non-sexual phase (73.6 µg/g) (U = 7, p < 0.0001), indicating that the EIA kits can reliably measure hormone metabolite titres (Fig. 1).

### 3.2 Markhor fP4M and fTM profiles and characterization of reproductive stages for both sexes

All five study animals (3F: 2M) were involved in reproductive activities. Analyses of annual fP4M and fTM data from male and female markhor individuals showed similar profiles, with peaks and baselines evident in both datasets. In females, despite differing baseline levels (Ivy = 204.92 µg/g, Sonia = 183.07 µg/g, and Supriya = 240.19 µg/g) (Table 3), fP4M levels increased in October from the baseline, remained elevated until April, indicating reproductive cyclicity during this period (Figure 2, Table 4). Consequently, fP4M levels returned to baseline in May and were maintained at a similar level until September, suggesting a non-reproductive phase. The reproductive phase (October-April) showed significantly higher fP4M titres than the non-reproductive phase (May-September) in all females (U = 492, p = 0.0001) (Tables 4 and 5). The gestation period was determined by sustained fP4M levels, detectable from October to April in all females, strongly indicating pregnancy (Tables 4 and 5), with a gestation of approximately seven months. However, fP4M levels varied throughout pregnancy. In the first 60 days, fP4M levels were lower (Ivy = 168-200 µg/g; Sonia = 640-690 µg/g; Supriya = 340-480 µg/g) (Table 5). In mid to late gestation, average fP4M levels increased significantly (Table 4), peaking near the end of pregnancy (Table 4). The pregnancy concluded with parturition, and fP4M levels sharply dropped to baseline from May onwards (Table 4). The female fTM baselines also varied across individuals Ivy = 117.02 µg/g, Sonia =135.75 µg/g, and Supriya = 191.89 µg/g) (Table 4). Except for Supriya, the other two females (Ivy and Sonia) showed no significant difference in fTM concentrations between the female reproductive and non-reproductive phases (U = 64, p = 0.0001). However, the overall fTM titre values were significantly different between the reproductive and non-reproductive phases (fTM: U = 1657, p = 0.001) (Table 4).

**Figure 2:**
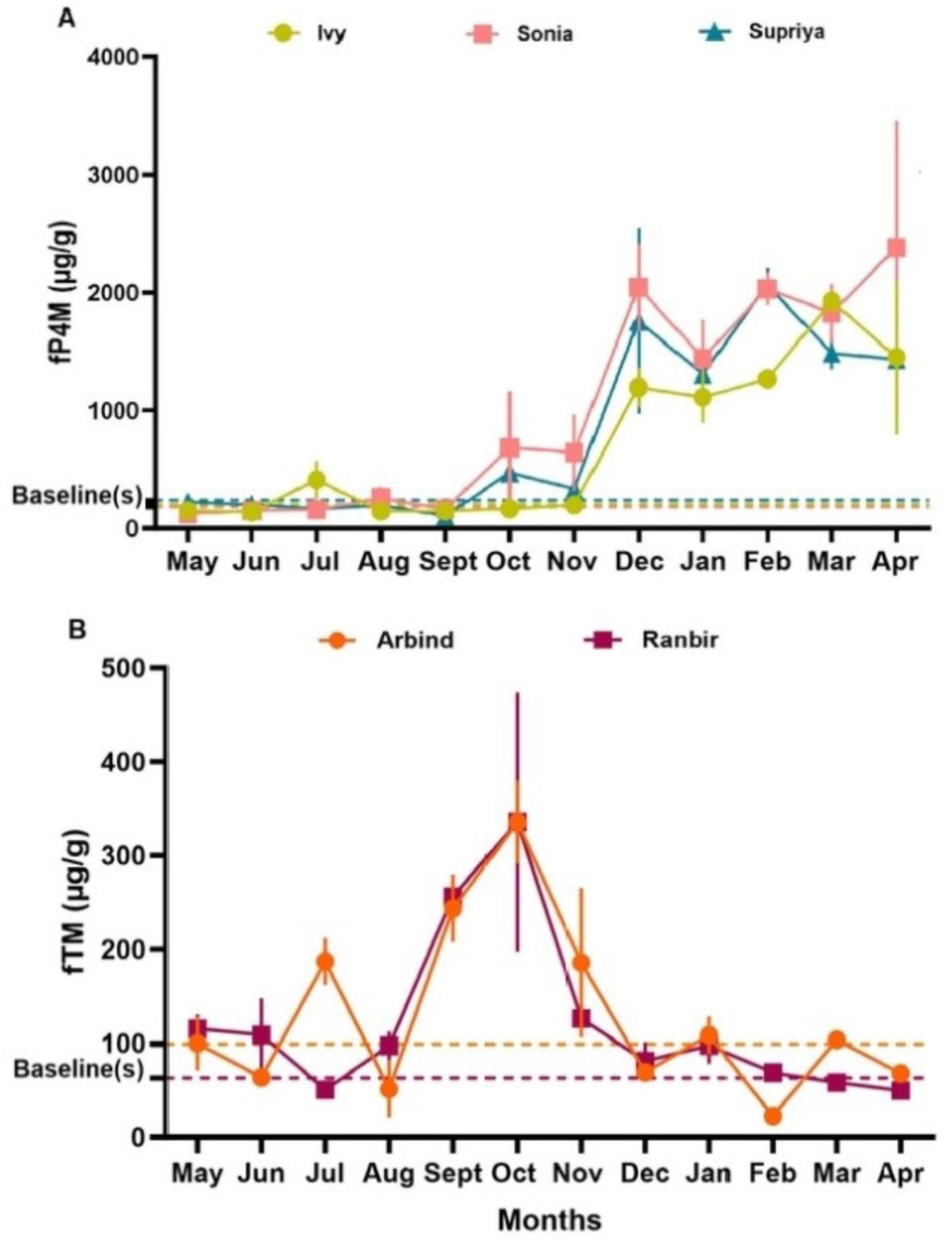
Annual reproductive profiles of male and female markhors. The top panel shows the fP4M profiles of all three study females, whereas the bottom panel shows the fTM profiles for the two males. The profiles are presented at monthly average levels. The dotted lines indicate the baselines derived for each individual.

**Table 3:** Measures of mean and baseline concentrations of fP4M and fTM across the study animals of PNHZP.

| Gender | Markhor name | Sample used | Fecal hormone | Overall mean conc. (M ± SEM) (μg/g) | Baseline conc. (μg/g) |
| --- | --- | --- | --- | --- | --- |
| ♀ | Ivy | 49 | Progesterone | 643.86 ± 95.43 | 204.92 |
| | | 48 | Testosterone | 211.52 ± 47.29 | 117.02 |
| | Sonia | 48 | Progesterone | 926.67 ± 143.38 | 183.07 |
| | | 47 | Testosterone | 210 ± 18.19 | 135.75 |
| | Supriya | 46 | Progesterone | 706.54 ± 116.69 | 240.19 |
| | | 45 | Testosterone | 305.36 ± 28.83 | 191.89 |
| ♂ | Arbind | 32 | Progesterone | 132.72 ± 7.3 | 170.86 |
| | | | Testosterone | 134.69 ± 18.42 | 99.21 |
| | Ranbir | 29 | Progesterone | 163.56 ± 30.45 | 161.09 |
| | | | Testosterone | 119.19 ± 16.49 | 63.4 |

**Table 4:**
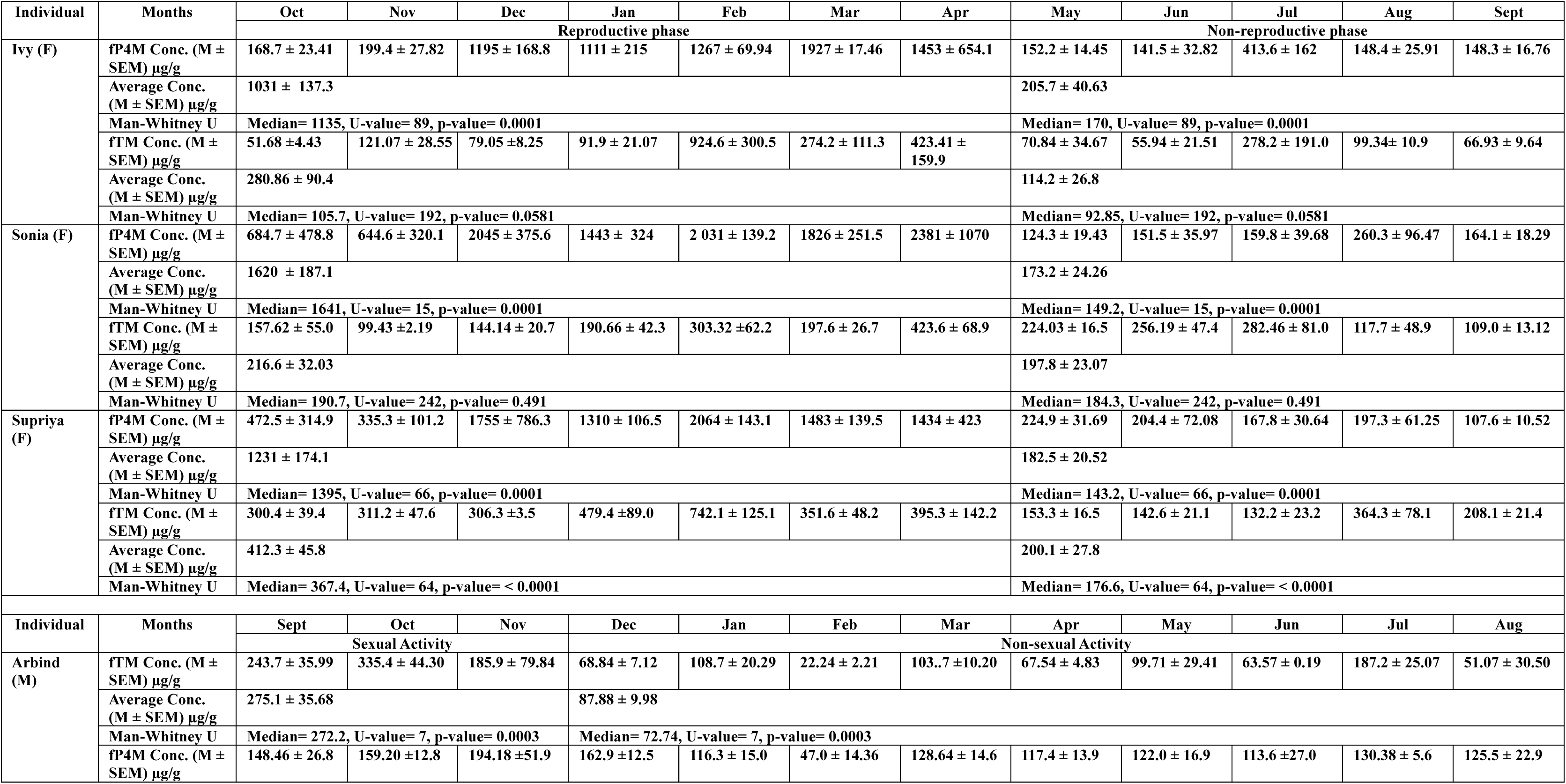
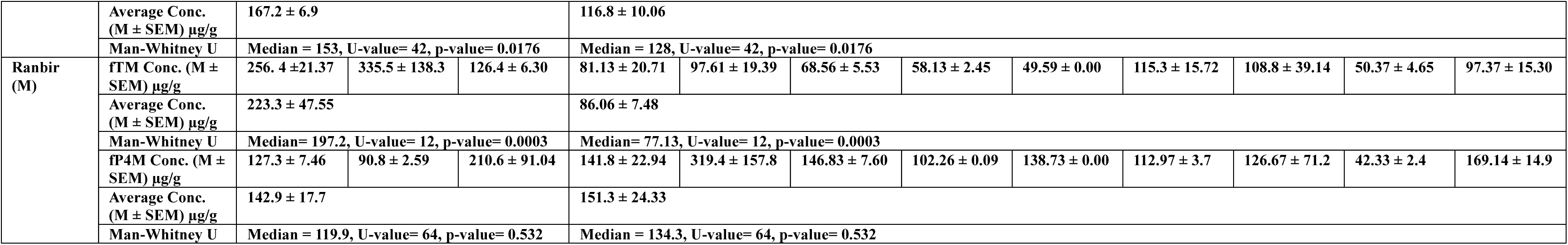
Measures of month-wise mean fP4M and fTM levels, and their comparative analyses between identified reproductive phases for all study animals.

**Table 5:** Comparative details of male and female markhor fP4M and fTM concentrations between identified reproductive phases.

| Target hormone | Comparison parameters | ♀ |  |  | ♂ |  |  |
| --- | --- | --- | --- | --- | --- | --- | --- |
|  |  | Reproductive Phase | Non-reproductive phase | Significance (p < 0.005) | Sexual Activity | Non-Sexual Activity | Significance (p < 0.005) |
| fP4M | Mean conc. (M ± SEM) $\mu\text{g/g}$ | 1292 ± 99.19 | 187.2 ± 17.02 | Yes | 159.4 ± 19.60 | 143.5 ± 18.87 | No |
|  | Man-Whitney U | Median=1340, U-value=492, p-value=0.0001 | Median=152.5, U-value=492, p-value=0.0001 |  | Median=141.7, U-value=273, p-value=0.2 | Median=132.7, U-value=273, p-value=0.2 |  |
| fTM | Mean conc. (M ± SEM) $\mu\text{g/g}$ | 299.9 ± 31.91 | 175.3 ± 19.53 | Yes | 250.9 ± 28.97 | 87.01 ± 6.25 | Yes |
|  | Man-Whitney U | Median=229.7, U-value=1657, p-value=0.001 | Median=133.7, U-value=1657, p-value=0.001 |  | Median=259.2, U-value=40, p-value=0.0001 | Median=73.51, U-value=40, p-value=0.0001 |  |

For each male, the onset of sexual activity was marked when the fTM concentrations exceeded the individual-specific baseline levels (Arbind = 170.86 µg/g; Ranbir = 161.09 µg/g) (Figure 2, Tables 3 and 4). Both males showed similar longitudinal hormone patterns, with fTM increasing in September and remaining elevated until November, before returning to baseline levels from December to August (Figure 2, Table 4). The comparison between the period of sexual activity (September-November) and non-sexual activity (December-August) was statistically significant (U=40, p=0.0001) (Table 4). The peaked fTM concentration was reflected in the expression of sexual behavior, such as copulation attempts that likely resulted in pregnancy, later confirmed by births in late April to early May. After establishing the baselines for fP4M (Arbind = 99.21 µg/g; Ranbir = 63.4 µg/g), both males displayed acyclic patterns, and there was no significant difference in fP4M concentrations between the sexual and non-sexual activity phases (U= 273, p=0.2) (Table 4).

All females showed significant differences in the monthly mean comparisons of fP4M concentrations (Ivy: χ2 = 34.42, p = 0.0003; Sonia: χ2 = 36.6, p = 0.0001; Supriya: χ2 = 28.82, p = 0.002), while the males did not show such a pattern (p = 0.16). However, significant differences in mean fTM concentrations across months were observed in both sexes (Arbind: χ2 = 25.75, p = 0.007; Ranbir: χ2 = 20.53, p = 0.038; Ivy: χ2 = 29.04, p = 0.002; Sonia: χ2 = 25.50, p = 0.007; Supriya: χ2 = 30.86, p = 0.001).

## Discussion

To the best of our knowledge, this study provides the first systematic non-invasive characterization of reproductive hormone profiles and stages (mating, gestation, and parturition) in both sexes of markhor. The reliable measurement of fecal progesterone and testosterone hormone metabolites was ensured through rigorous EIA validation, including parallelism, accuracy, and biological validations in markhor. The inter- and intra-assay variations were ≤ 10% for both target hormones, indicating reliable technical reproducibility and minimizing errors from multiple pipetting and handling (Sink et al., 2008). Therefore, such a fecal-based approach, combined with long-term spatio-temporal sampling, can now be used to support comprehensive monitoring of reproductive activity in both male and female markhor individuals in the wild and in managed care, thereby contributing to improved animal welfare, management, and conservation of this species (Peel et al., 2005; Schwarzenberger, 2007).

The EIA analysis of P4 hormone in female markhor feces distinguished between pregnant and non-pregnant phases. The endocrine assessment of fP4M proved effective, as P4 levels during pregnancy were consistently significantly higher than in the non-pregnant phase across all three females observed during the study. Elevated progesterone levels were observed in all female markhors from October to April, indicating pregnancy. Elevated progesterone levels are known to be associated with corpus luteum formation and its maintenance throughout pregnancy, leading to increased progesterone levels (Swanson et al., 1995; Verhage et al., 1976). Additionally, progesterone is crucial in promoting sexual receptivity, working together with other sex steroids (Witt et al., 1994). In this case, the increase in fP4M from October onwards indicates ovarian function following mating, leading to pregnancy in subsequent months. Such elevated P4 levels for ∼212 days (October to April, ∼7 months) suggest the gestation period of the markhor (Fig. 3). The timing of parturition observed in this study largely corroborates the available information on a gestation period of ∼155 days (Menon, 2003), whereas field observations indicate that birth occurs in April in captivity, and May in the wild (Menon, 2003) (Fig. 3). Pre-birth, a significant decline in circulating fP4M levels was observed in all the females at the zoo, a pattern regularly reported in other mammalian species (Short, 1960). The non-pregnant phase, from May to September, exhibited relatively low progesterone levels (below baseline), underscoring P4’s essential role in stabilizing and preparing the endometrium before implantation and in regulating ovulation (Jabbour et al., 2006). Overall, our current study emphasizes the importance of independently validating fecal progestogens alongside measures of pregnancy, and the EIA successfully distinguished the pregnant state from the non-pregnant state using fP4M as an endocrine biomarker. However, the overall fP4M concentrations in males did not significantly differ between the sexual activity and non-sexual activity phases. The fP4M titre was approximately 1.07 times higher during the sexual activity period, indicating the potential role of progesterone in androgen-dependent sexual behavior in males, as observed in other mammals (Witt et al., 1994).

**Figure 3:**
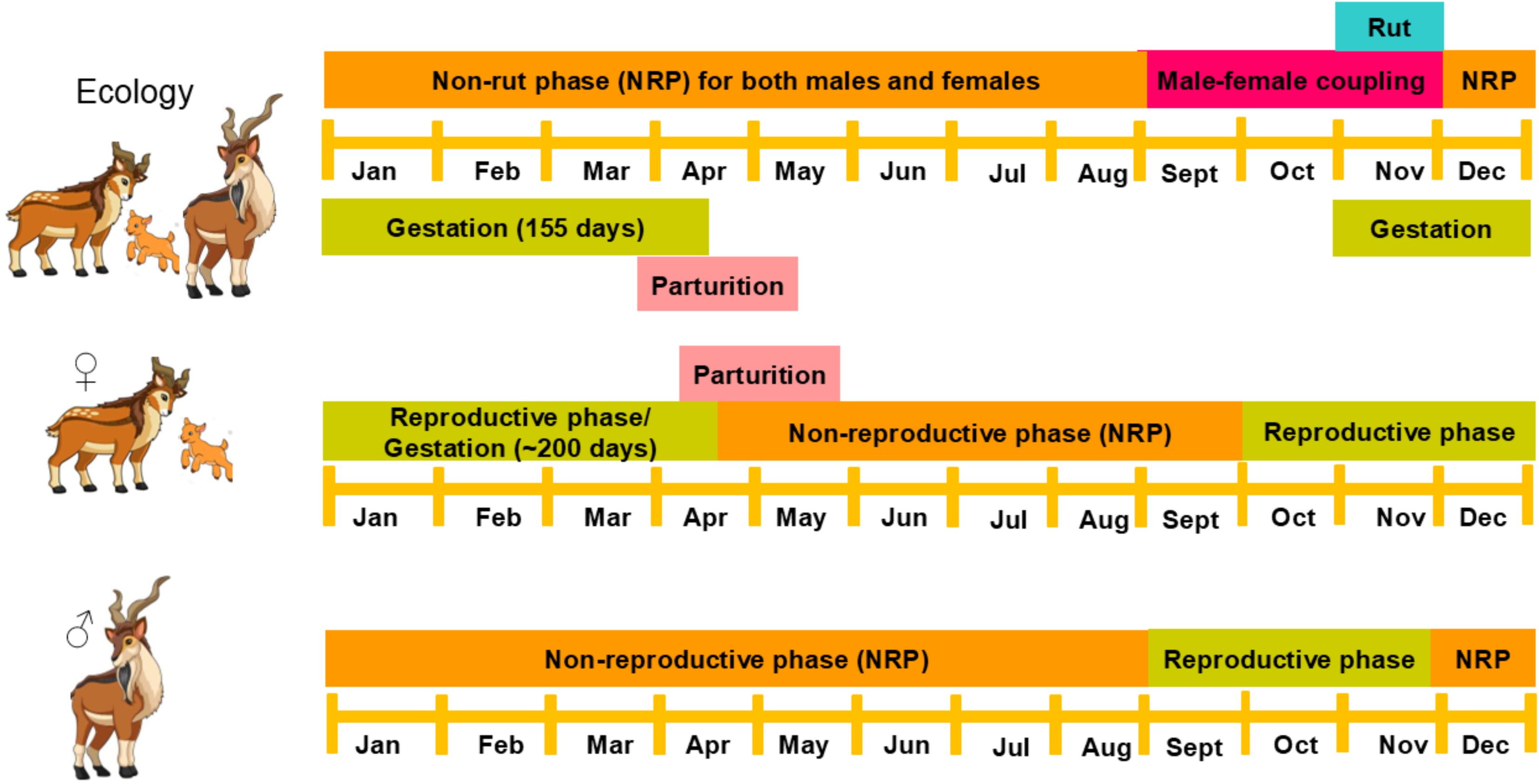
Comparative graphical depiction of the available ecological and endocrinological findings for both male and female markhor. The endocrinological assessments of the reproductive phases largely align with field observations.

The testosterone (fTM) concentrations fluctuated between periods of sexual activity and inactivity, with significantly higher levels during September to November and below baseline from December to August, respectively, in male markhor individuals. Androgens, such as testosterone, are key indicators of sexual maturity that influence the development of the male reproductive tract, induce spermatogenesis, and affect sexual behavior and libido in many mammalian species (McQuaid and Tanrikut, 2014; Jewgenow and Songsasen, 2014; Holt et al., 2014; Lanyon and Burgess, 2014). Testosterone secretion before the breeding season is essential for completing the spermatogenic cycle and transporting sperm through the epididymis (Gomes, 1978). This pattern of higher testosterone levels is supported by the earlier markhor study (Bezjian et al., 2013), where males exhibited seasonality, and the concentration of fecal testosterone peaked in November. The seasonal breeding behavior was supported by the synchronous progesterone peaks observed in female markhor, which coincided with increased testosterone levels in males. The timing of heightened fTM concentrations matches visual ecological records, in which male-female pairings were also observed between September and November, during the rut or breeding phase in this species (Menon, 2003) (Fig. 3). Lower fecal testosterone from December to August indicates a period of reproductive decline and a potential infancy of spermatozoa, a pattern observed in seasonal breeding species (Holt et al., 2014). The quantification of fecal T biomarkers using EIA produced similar results, as hormonal data complemented ecological studies and confirmed that the non-rut phase in markhor lasts from December to August (Menon, 2003). Interestingly, females exhibited high fTM titres during the reproductive or cycling phase (October-April). The presence of testosterone during this period indicates that testosterone plays a key role in female sexual behavior, alongside maintaining cognition, metabolism, and mediating directly to estradiol via aromatization (Cipriani et al., 2021; Davis and Wahlin-Jacobsen, 2015). These detectable testosterone levels also suggest that testosterone may play a key role in regulating secondary sexual characteristics in markhor females, as observed in other vertebrates. (Goymann and Wingfield, 2014; Staub and De Beer, 1997). Taken together, the non-invasive measurement of both reproductive hormones accurately reflected ovarian function in females, assisting in establishing the timeline for mating, gestation length, and parturition in female markhors, as well as the breeding season timeline in males. Therefore, such approaches can significantly support the husbandry and breeding management of this species by determining the optimal pairing period for breeding, diagnosing pregnancy, and predicting parturition in both captive and wild populations.

The systematic monitoring of reproductive hormones also revealed individual variation in hormone levels in both sexes. All individuals (3F: 2M) showed sex-specific synchronized patterns in hormone titres, but their hormone levels differed among them. This has much larger implications in their large-scale use in markhor physiology. In mammalian systems, reproductive hormone titres may vary due to intrinsic (individual health status, stress levels, etc.) and extrinsic (environment, diet, welfare conditions, etc.) factors, but the cyclicity of the reproductive phases is expected to be synchronous (Asher et al., 1999; Devra et al., 2010). Our data also confirmed this pattern, and therefore, the outcomes can be readily applied to reproductive monitoring in the wild and in other captive facilities. If suitably applied, such tools have tremendous potential for monitoring the populations of other endangered species across high-altitude landscapes worldwide.

## Acknowledgement

We thank the Principal Chief Conservator of Forest (PCCF) and the Chief Wildlife Warden (CWLW), West Bengal, and the Member Secretary of the West Bengal Zoo Authority (WBZA) for their support for the Study. We are also grateful to the various teams at PNHZP, especially the veterinary, research, and animal-keeping teams, for their cooperation and valuable insights throughout the Study. Our sincere thanks go to the Zoo’s administrative staff for their cooperation and ongoing support with the required paperwork. We acknowledge the assistance of Ms. Nabanita Ghosh, Dr. Barkha Subba, Miss Prishka Pariyar, Miss Rohini Chetri, and Miss Pranita Subba during the project. We also thank the Director, Dean, and Research Coordinator, Wildlife Institute of India for their support. Funding for this work was provided by PNHZP, supported by the West Bengal Zoo Authority.

## Supplementary Figure

**Supplementary Figure 1:**
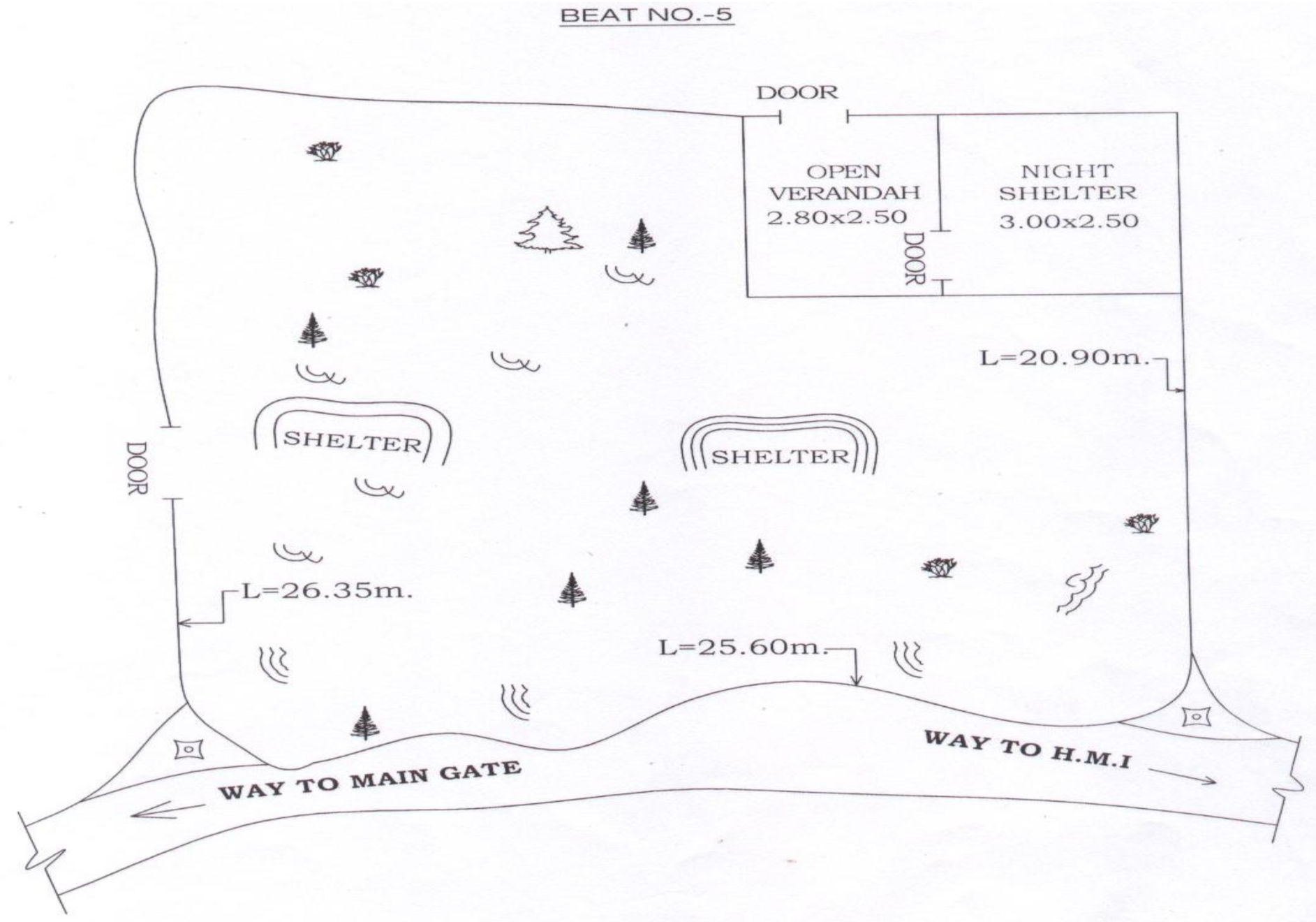
Schematic representation of the markhor enclosure at PNHZP.

## Supplementary Table

**Supplementary Table 1:** Daily feed details of Markhor at PNHZP.

| SI No | Feed items | Quantity provided per animal | Feed timings |
| --- | --- | --- | --- |
| 1 | Crushed maize | 250 gms | 9:00 am |
| 2 | Gram | 70 gms |  |
| 3 | Crushed wheat | 60 gms |  |
| 4 | Barley | 120gms |  |
| 5 | Wheat bran | 140 gms |  |
| 6 | Salt | 5 gms |  |
| 7 | Pulses (moong/ masoor) | 40 gms |  |
| 8 | Molasses | 50gms |  |
| 9 | Turmeric | 5 gms |  |
| 10 | Green fodder | 8 kgs | 4 kgs at 9:00 am<br>4 kgs at 4:00 pm |

**Supplementary Table 2:**
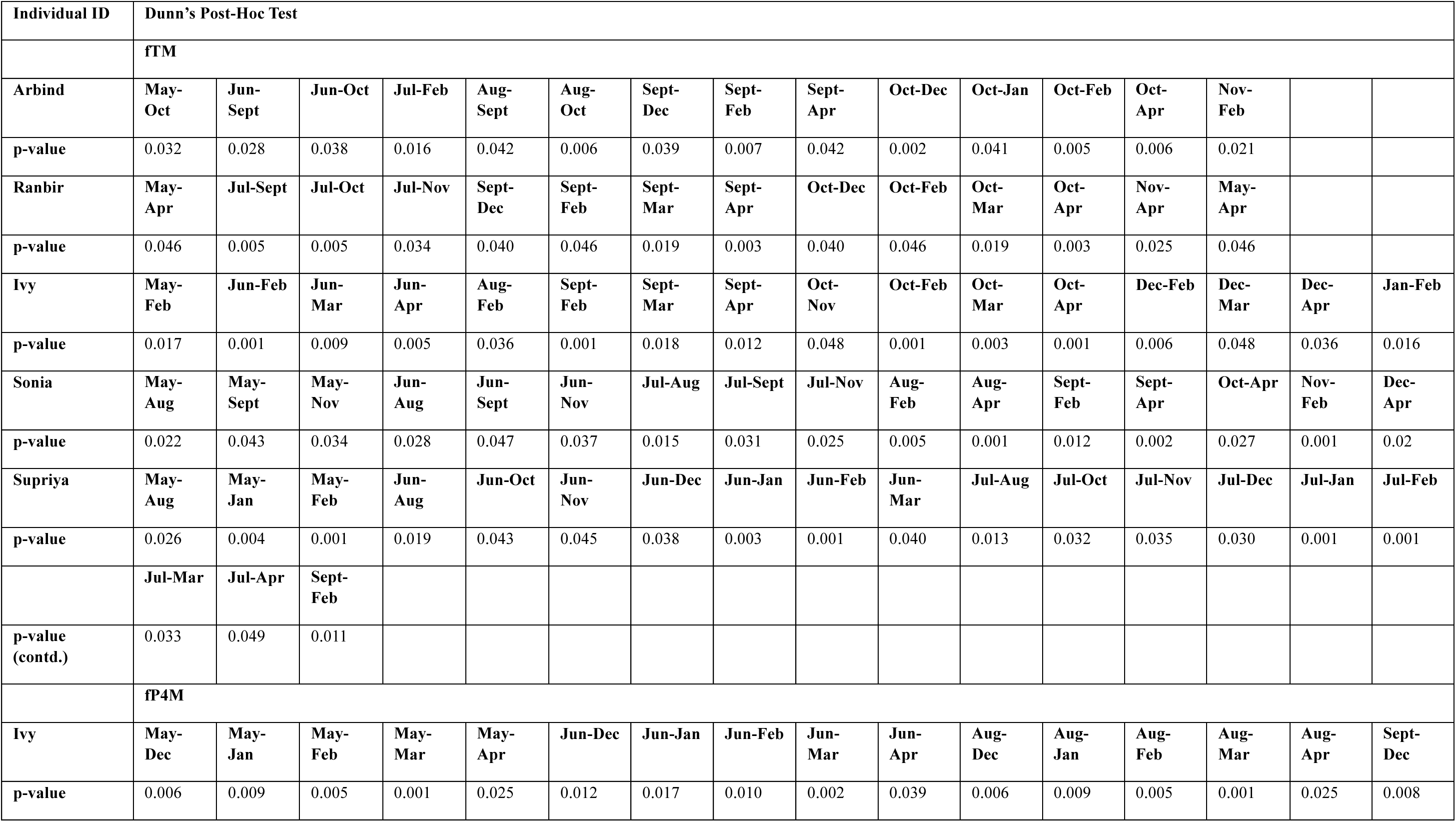
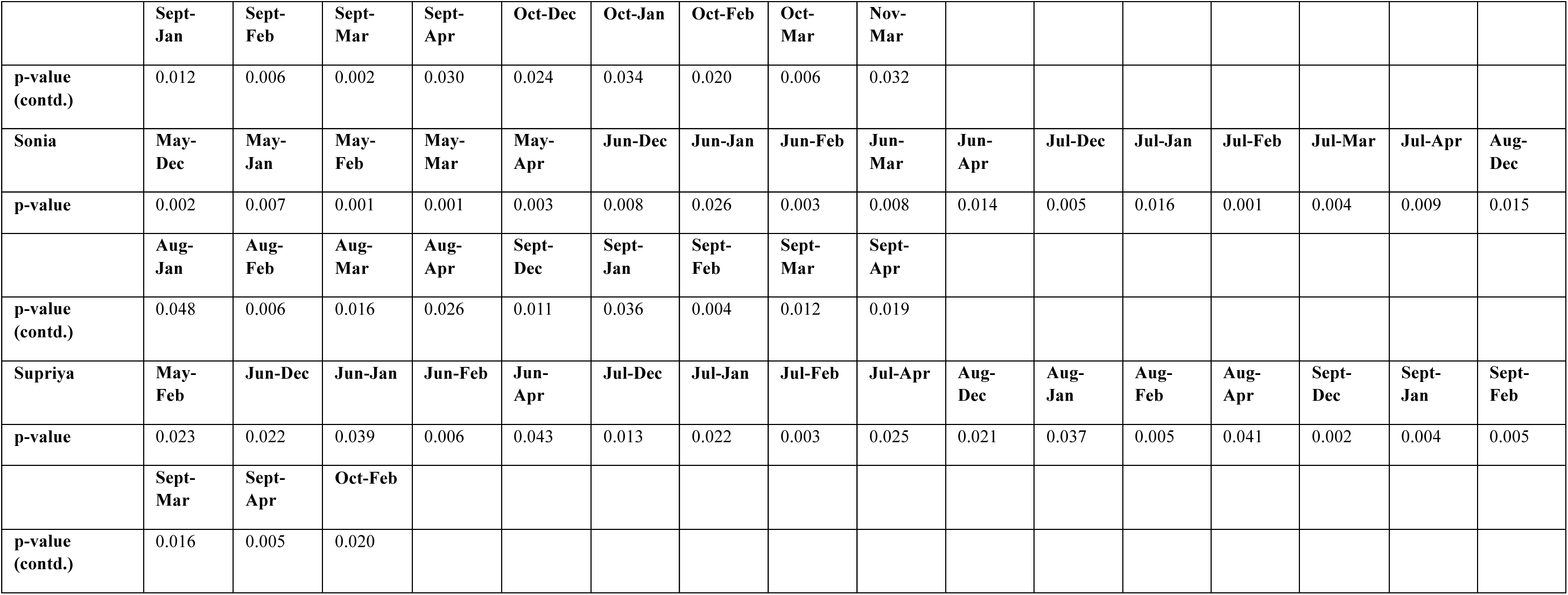
Comparative analyses of month-wise median concentrations of fP4M and fTM measures in all study subjects. The Dunn’s post-hoc comparisons reveal multiple significant (p-value < 0.005) differences.

| Individual ID | Dunn's Post-Hoc Test |  |  |  |  |  |  |  |  |  |  |  |  |  |  |  |
| --- | --- | --- | --- | --- | --- | --- | --- | --- | --- | --- | --- | --- | --- | --- | --- | --- |
|  | fTM |  |  |  |  |  |  |  |  |  |  |  |  |  |  |  |
| Arbind | May-Oct | Jun-Sept | Jun-Oct | Jul-Feb | Aug-Sept | Aug-Oct | Sept-Dec | Sept-Feb | Sept-Apr | Oct-Dec | Oct-Jan | Oct-Feb | Oct-Apr | Nov-Feb |  |  |
| p-value | 0.032 | 0.028 | 0.038 | 0.016 | 0.042 | 0.006 | 0.039 | 0.007 | 0.042 | 0.002 | 0.041 | 0.005 | 0.006 | 0.021 |  |  |
| Ranbir | May-Apr | Jul-Sept | Jul-Oct | Jul-Nov | Sept-Dec | Sept-Feb | Sept-Mar | Sept-Apr | Oct-Dec | Oct-Feb | Oct-Mar | Oct-Apr | Nov-Apr | May-Apr |  |  |
| p-value | 0.046 | 0.005 | 0.005 | 0.034 | 0.040 | 0.046 | 0.019 | 0.003 | 0.040 | 0.046 | 0.019 | 0.003 | 0.025 | 0.046 |  |  |
| Ivy | May-Feb | Jun-Feb | Jun-Mar | Jun-Apr | Aug-Feb | Sept-Feb | Sept-Mar | Sept-Apr | Oct-Nov | Oct-Feb | Oct-Mar | Oct-Apr | Dec-Feb | Dec-Mar | Dec-Apr | Jan-Feb |
| p-value | 0.017 | 0.001 | 0.009 | 0.005 | 0.036 | 0.001 | 0.018 | 0.012 | 0.048 | 0.001 | 0.003 | 0.001 | 0.006 | 0.048 | 0.036 | 0.016 |
| Sonia | May-Aug | May-Sept | May-Nov | Jun-Aug | Jun-Sept | Jun-Nov | Jul-Aug | Jul-Sept | Jul-Nov | Aug-Feb | Aug-Apr | Sept-Feb | Sept-Apr | Oct-Apr | Nov-Feb | Dec-Apr |
| p-value | 0.022 | 0.043 | 0.034 | 0.028 | 0.047 | 0.037 | 0.015 | 0.031 | 0.025 | 0.005 | 0.001 | 0.012 | 0.002 | 0.027 | 0.001 | 0.02 |
| Supriya | May-Aug | May-Jan | May-Feb | Jun-Aug | Jun-Oct | Jun-Nov | Jun-Dec | Jun-Jan | Jun-Feb | Jun-Mar | Jul-Aug | Jul-Oct | Jul-Nov | Jul-Dec | Jul-Jan | Jul-Feb |
| p-value | 0.026 | 0.004 | 0.001 | 0.019 | 0.043 | 0.045 | 0.038 | 0.003 | 0.001 | 0.040 | 0.013 | 0.032 | 0.035 | 0.030 | 0.001 | 0.001 |
|  | Jul-Mar | Jul-Apr | Sept-Feb |  |  |  |  |  |  |  |  |  |  |  |  |  |
| p-value (contd.) | 0.033 | 0.049 | 0.011 |  |  |  |  |  |  |  |  |  |  |  |  |  |
|  | fP4M |  |  |  |  |  |  |  |  |  |  |  |  |  |  |  |
| Ivy | May-Dec | May-Jan | May-Feb | May-Mar | May-Apr | Jun-Dec | Jun-Jan | Jun-Feb | Jun-Mar | Jun-Apr | Aug-Dec | Aug-Jan | Aug-Feb | Aug-Mar | Aug-Apr | Sept-Dec |
| p-value | 0.006 | 0.009 | 0.005 | 0.001 | 0.025 | 0.012 | 0.017 | 0.010 | 0.002 | 0.039 | 0.006 | 0.009 | 0.005 | 0.001 | 0.025 | 0.008 |
|  | **Sept-Jan** | **Sept-Feb** | **Sept-Mar** | **Sept-Apr** | **Oct-Dec** | **Oct-Jan** | **Oct-Feb** | **Oct-Mar** | **Nov-Mar** |  |  |  |  |  |  |  |
| **p-value (contd.)** | 0.012 | 0.006 | 0.002 | 0.030 | 0.024 | 0.034 | 0.020 | 0.006 | 0.032 |  |  |  |  |  |  |  |
| **Sonia** | **May-Dec** | **May-Jan** | **May-Feb** | **May-Mar** | **May-Apr** | **Jun-Dec** | **Jun-Jan** | **Jun-Feb** | **Jun-Mar** | **Jun-Apr** | **Jul-Dec** | **Jul-Jan** | **Jul-Feb** | **Jul-Mar** | **Jul-Apr** | **Aug-Dec** |
| **p-value** | 0.002 | 0.007 | 0.001 | 0.001 | 0.003 | 0.008 | 0.026 | 0.003 | 0.008 | 0.014 | 0.005 | 0.016 | 0.001 | 0.004 | 0.009 | 0.015 |
|  | **Aug-Jan** | **Aug-Feb** | **Aug-Mar** | **Aug-Apr** | **Sept-Dec** | **Sept-Jan** | **Sept-Feb** | **Sept-Mar** | **Sept-Apr** |  |  |  |  |  |  |  |
| **p-value (contd.)** | 0.048 | 0.006 | 0.016 | 0.026 | 0.011 | 0.036 | 0.004 | 0.012 | 0.019 |  |  |  |  |  |  |  |
| **Supriya** | **May-Feb** | **Jun-Dec** | **Jun-Jan** | **Jun-Feb** | **Jun-Apr** | **Jul-Dec** | **Jul-Jan** | **Jul-Feb** | **Jul-Apr** | **Aug-Dec** | **Aug-Jan** | **Aug-Feb** | **Aug-Apr** | **Sept-Dec** | **Sept-Jan** | **Sept-Feb** |
| **p-value** | 0.023 | 0.022 | 0.039 | 0.006 | 0.043 | 0.013 | 0.022 | 0.003 | 0.025 | 0.021 | 0.037 | 0.005 | 0.041 | 0.002 | 0.004 | 0.005 |
|  | **Sept-Mar** | **Sept-Apr** | **Oct-Feb** |  |  |  |  |  |  |  |  |  |  |  |  |  |
| **p-value (contd.)** | 0.016 | 0.005 | 0.020 |  |  |  |  |  |  |  |  |  |  |  |  |  |

